# Discovery of Novel R-Selective Aminotransferase Motifs through Computational Screening

**DOI:** 10.1101/2024.08.21.608959

**Authors:** Ashish Runthala, Pulla Sai Satya Sri, Aayush S Nair, Murali Krishna Puttagunta, T Chandra Sekhar Rao, Vajrala Sreya, Ganugapati Reshma Sowmya, Koteshwara Reddy G

## Abstract

Transaminases, enzymes facilitating amino group transfers, are divided into four subfamilies: D-alanine transaminase (DATA), L-selective Branched chain aminotransferase (BCAT), 4-amino-4-deoxychorismate lyase (ADCL), and R-selective aminotransferase (RATA). RATA enzymes are particularly valuable in biocatalysis for synthesizing chiral amines and resolving racemic mixtures, yet their identification in sequence databases is challenging due to the lack of robust motif-based screening methods. By constructing a transaminase sequence dataset and categorizing them into subfamilies, we re-screened conserved motifs and explored novel ones. Phylogenetic clustering and structural localization of these motifs on Alphafold-predicted protein models validated their importance. For ADCL, BCAT, DATA, and RATA datasets, we discovered 5, 7, 10, and 2 novel motifs, respectively. Additionally, unique residue patterns were identified, underscoring their structural significance. This motif-based computational approach promises to unveil novel RATA enzymes for biocatalytic applications.

## Introduction

Transaminases or aminotransferases are present in all the living organisms, and catalyze the reaction between an α-keto acid and an amino acid (Lilly et al., 2006). Transamination reactions, specific to amino and keto groups, play a vital role within tissues, their directionality dictated by reactant concentrations. This reversibility and stereoselective nature of the reactions is used in synthetic chemistry applications to synthesize the valuable chiral amines (Dunathan et al. 1968; Faber & Kroutil, 2005; Schiroli & Peracchi, 2015). These enzymes are named after their reactants, such as glutamic-pyruvic transaminase or GPT, which catalyzes the reaction between glutamic acid and pyruvic acid to form alpha ketoglutaric acid and alanine (Jenkins et al., 1957). Being biologically important to drive the synthesis of amino acids, transamination reaction deaminates an amino acid or NH_2_-donor amine, and aminates the NH_2_-acceptor aldehyde/ketone/keto acid, synthesizing a new amine or amino acid (Guo et al., 2017). The vitamin B6-derived cofactor, pyridoxal 5’-phosphate (PLP), binds to the enzyme’s conserved catalytic lysine via an imine bond, functioning as an electron sink and stabilizing the reaction intermediates (Shimizu et al., 2010). Transaminases play a vital role in numerous metabolic processes (Rudat et al. 2012; Lieu et at. 2020). Their analysis is essential for various purposes, including assessing liver function, diagnosing and monitoring liver diseases, understanding metabolic pathways (Jamithireddy et al. 2020; Vulichi et al. 2020), evaluating drug safety, and investigating genetic disorders and biochemical synthesis of drugs (Kaplan, 1986; Dufour et al. 2000; Pratt & Kaplan, 2000; Schuppan, & Afdhal, 2008; Senior, 2014). This comprehensive understanding of transaminases is crucial for both clinical practice and medical research. However, identifying new, functionally efficient aminotransferase sequences from protein databases remains a challenge, despite its necessity.

Based on the functional or evolutionary relationships, transaminases are classified under seven major structural groups (Slabu et al., 2017). Protein Family Databases (PFAM) further groups these enzymes into 6 subfamilies as per their common fold architecture and functional as well as evolutionary relationships (Buss et al., 2018). Stereoselective S and R-transaminases with minimal sequence similarity belong to distinct Fold I and IV groups (Jansonius, 1998; Percudani & Peracchi, 2009), identified by PFAM through the conserved sequence motifs (van Oosterwijk et al., 2016). These enzymes transfer the amino group from a donor amine substrate to an amine acceptor substrate, which is usually a ketone compound. While the Fold-I encompasses L/S-selective α-TA enzymes and none of its enzymes exhibits a D/R-selection, the Fold-IV set engulfs the D-as well as L-selective enzymes, and is a cluster of both α-TA and ω-TA, with four major classifications: D-alanine transaminase (DATA) (Jia et al., 2022), L-selective Branched chain aminotransferase (BCAT) (Ichihara et al., 1966), and 4-amino-4-deoxychorismate lyase (ADCL) (Jhee et al., 2000) and very rare R-selective aminotransferase (RATA) (Lyskowski et al., 2014) sequences.

Aminotransferases exhibit substantial sequence diversity, even among enzymes catalyzing similar reactions (Mehta et al., 1993), making it difficult to define the universally conserved motifs across the four classes, and complicating the sequence search algorithm. Additionally, many structural motifs associated with aminotransferases, such as the pyridoxal phosphate binding site, are shared across different enzyme families (Mehta et al., 2000), confounding the motif-based discrimination of sequences. Further complicating matters, the functional divergence within the aminotransferase superfamily is not always reflected in the primary sequence, and subfamilial members have been shown to have a very low sequence similarity (Grishin et al., 1995). Protein sequence databases may also lack comprehensive representation of aminotransferase diversity, as newly discovered or understudied enzymes may lack well-defined conserved motifs (Mistry et al. 2021). To overcome these limitations, a strategic screening of the conserved, yet unknown, motifs need to be implemented by integrating the robustly-scoring profile-based searches (Söding, 2005) for the protein sequences. Ultimately, a combination of computational and experimental validation may be necessary to reliably identify and confirm aminotransferases within large protein datasets. Though significant research has been done to search and industrially tailor the transaminases for improved functions, minimal research has been done to decode the novel motifs, although it still holds the major crux of the entire research done so far, and is current research focus to discover novel enzymes exhibiting the improved properties.

## Materials and Methods

### Building the sequence dataset

For building the sequence dataset for the downstream analysis, the HMMER database (Potter et al., 2018) is used to screen the topranked sequences for the engineered transaminase (3WWJ) and the native protein (3WWH). For increasing the algorithmic accuracy, the redundant entries, functionally different and partial protein sequences are then discarded to result in constructing the desired dataset. Further, as per the sequence length of the reduced dataset of experimentally resolved transaminase structures, a threshold of 200-400 residues is deployed to search the functionally correct set of transaminases, as shown as the initial step of the methodology, represented in Figure1.

**Figure 1:**
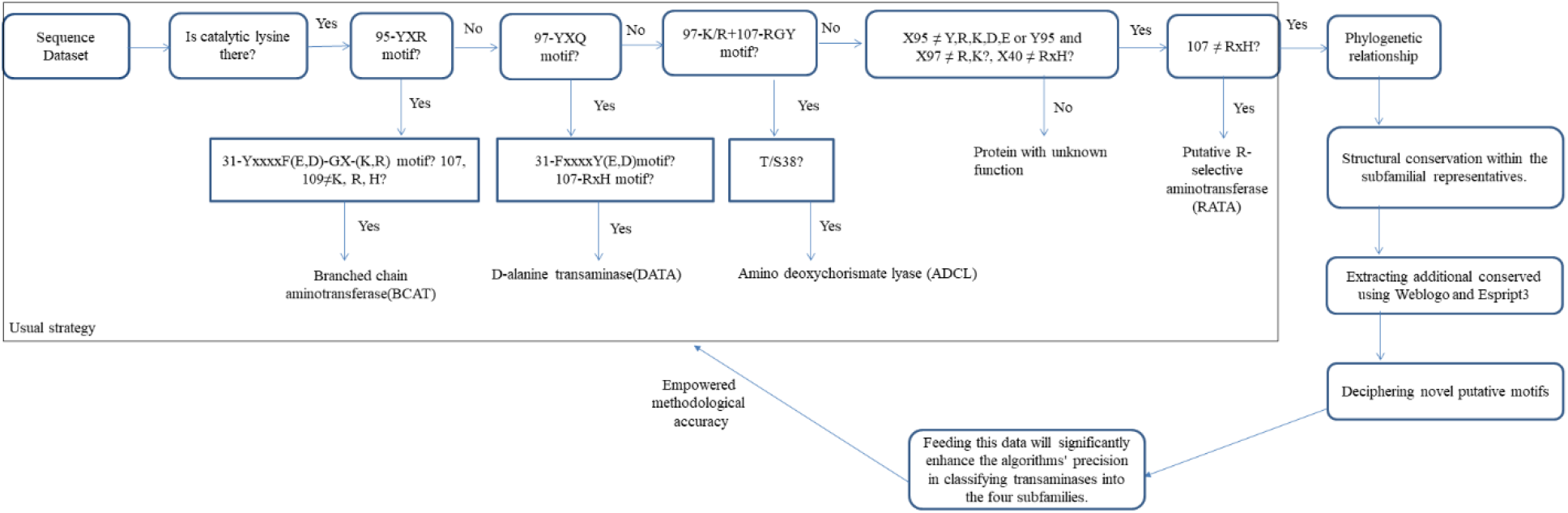
Computational methodology used for the analysis of the transaminase proteins.

### Building the subfamilial subsets

As per Hohne’s protocol (Hohne et al., 2010), the retrieved sequences are categorized into subfamilies. To reliably do the scrutiny and extract the true RATA sequences lastly, the non-redundant functionally known protein structures are extracted, and are aligned with the respective sequence subsets. For constructing a more accurate alignment, the hidden markov model (HMM) based Clustal omega protocol is utilized through the HHPred server (Söding et al., 2005). As per the conserved and functionally crucial signature segments, the sequence datasets are reliably categorized into the defined subfamilies, i.e. ADCL, BCAT, DATA, RATA, and the leftover entries with atleast two functionally different motifs are categorized as the proteins with unidentified function (PUF).

### Estimating the evolutionary and conformational diversity

To infer the relevant phylogenetic linkages within the chosen set of sequences/species, statistically significant evolutionary relationships are usually taken into consideration. Since an exact evolutionary relationship can only be extracted with a precise sequence alignment, the sequence dataset of all the subfamilies is aligned using the hidden Markov model-based clustal-omega module (Söding et al., 2005). To derive the evolutionary meaningful relationships between the sequences, the constructed alignment is fed to IQ-tree (Trifinopoulos et al., 2016) for building their reasonable phylogenetic solution on the basis of maximum likelihood methodology. To enhance the algorithmic accuracy, 10,000 bootstrap alignments are constructed using the ultrafast methodology, and iterating over 10,000 iterations, the optimal phylogenetic placement of every sequence is established using a minimum correlation coefficient of 0.99. Lastly, the consensus-derived phylogenetic tree model is viewed and analyzed using ITOL (Letunic et al., 2007) to infer the relationships.

To estimate the structural conservation across the catalytically crucial substructures, the longest sequence of each subfamilial subset is considered as the representative entry (Steinegger & Söding, 2018). The tertiary structures of the 4 key sequences are then predicted using the deep-learning based modelling methodology AlphaFold (Senior et al., 2020) to enhance the credibility of the downstream evolutionary analysis. The conserved core is then analyzed for the representatives using Chimera, and their alignment is plotted onto the representative RATA protein using Espript3 (Bonneau et al., 2009). It would allow us to map the extent of sequence and topological similarity across these representatives.

### Analyzing the functionally crucial residues of the four subfamilies

A functionally crucial residue should be maximally conserved in the subfamily, and hence, it should form a basis to discriminate between the subfamilies (Bezsudnova et al., 2020). Hence, rather than only focusing on the substantially conserved/variant residue positions, the overall network of conservation should be evaluated to properly decode the meaningful regions, mandatorily required to govern the function. With special consideration to the promiscuity of the active-site, the evaluated region should maximally encompass the key active site residues and the catalytic loci that initiate/assist the functional regions.

Hence, the HMM-based sequence alignments of various subfamilies are fed to Weblogo3 (Crooks et al., 2022) to assess the degree of sequence conservation and decode the discriminatory centers. Each logo consists of symbol stacks for each sequence position, where the stack height shows sequence conservation and the symbol each amino acid at that position (Crooks et al., 2022).

## RESULTS AND DISCUSSION

### Building the sequence dataset

Screening HMMER for 3WWH and 3WWJ proteins, a set of 1035 sequences is constructed, of which only the redundant entries, functionally different and partial height within the stack reflects the relative frequency of protein sequences are then discarded to result into the reduced dataset of 543 proteins. However, as the sequence length of experimentally solved protein structures ranges from 200-400 residues, the constructed set is again filtered, yielding 247 sequences, as illustrated in the following Figure2.

**Figure 2:**
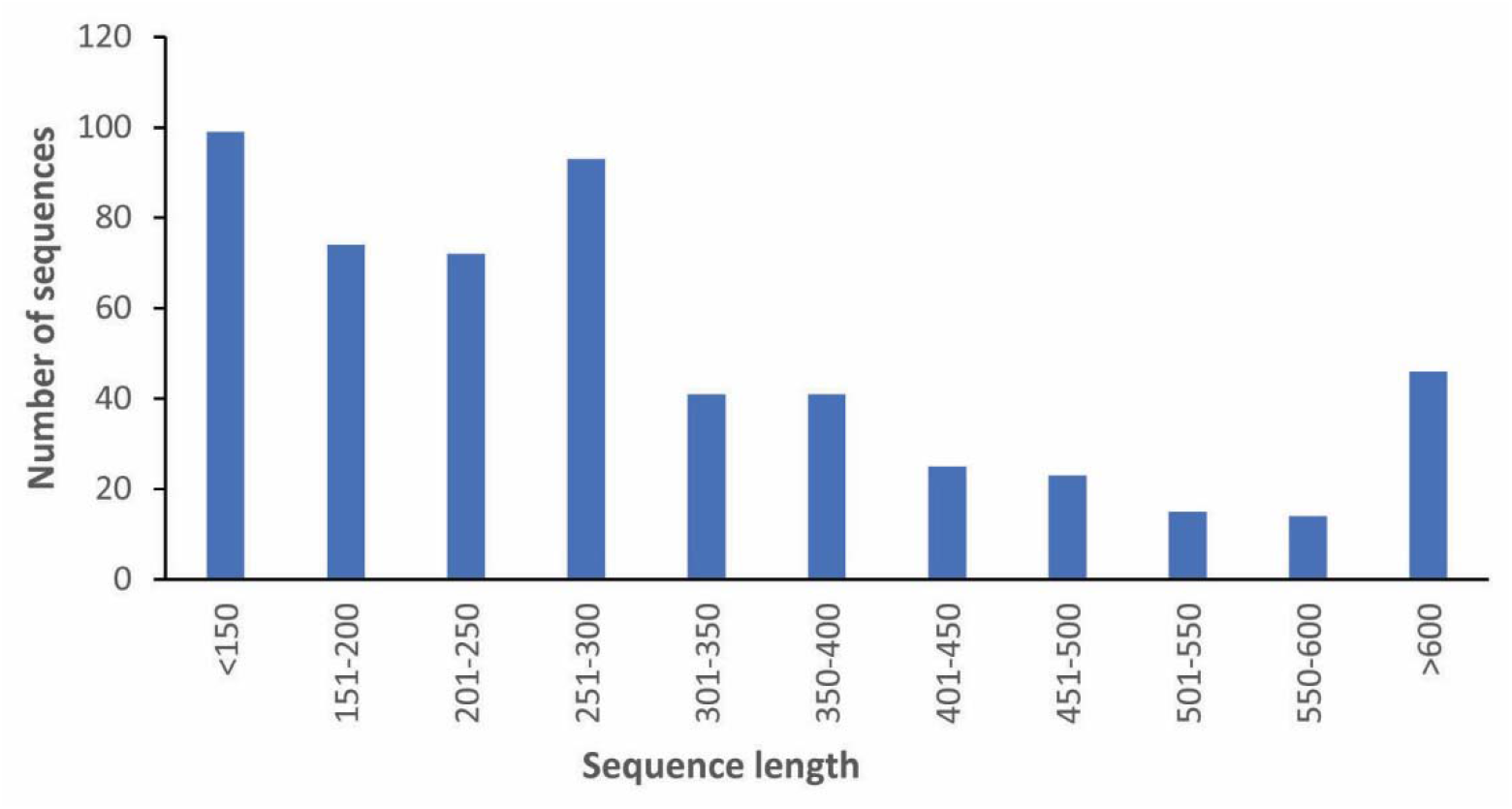
Sequence length clusters of the sequence dataset, indicating that Omega-transaminase proteins majorly encode 200-400 residues.

### Building the subfamilial subsets

Based on the catalytic lysine, mapped as per the experimentally resolved structures, the selected sequences are categorized into transaminases and functionally different sequences, as represented in Figure1. This scrutiny reliably filtered out a set of 63 transaminases, and allowed us purge the meaningless proteins, not having the catalytic lysine. The transaminases are then grouped into the subfamilies as per their conserved motifs. For classifying the entries into subfamilies, the pre-reported conserved signatures are used (Hohne et al., 2010). If at 95th position YXR motif is present and at 107 and 109 positions amino acids K, R, H are not present if these conditions are satisfied the sequence is considered as BCAT classification (Hirotsu et al., 2005). If at 97th position YXQ motif is present and at 107 position RH motif is present, if the condition is satisfied the sequence is considered as DATA classification. If at 97th position amino acids K/R is present, at 38th position amino acids S/T is present and at 107 position RGY motif is present, if these conditions are satisfied the sequence is considered as ADCL classification (Bezsudnova et al. 2019). If at 107th position amino acids R/H is not present, if the condition is satisfied the sequence is considered as RATA, otherwise the entry is considered as protein sequence with unknown function (PUF), left undiscussed in the earlier paper (Hohne et al., 2010). The analysis thus yielded a set of 9, 19, 5, 12, and 18 ADCL, BCAT, DATA, RATA, and PUF proteins, respectively.

### Estimating the evolutionary and conformational diversity

Constructing the sequence alignment using HMM-Clustal Omega (Söding et al., 2005) and building the phylogenetic tree with IQ-Tree (Trifinopoulos et al., 2016), based on 10,000 iterations of 10,000 bootstrap alignments with a correlation coefficient score of 0.99, is expected to enhance algorithmic accuracy. Reliably integrating the associated subfamilial entries to show the consistently resolved branch lengths, the methodology yielded the phylogenetic tree with a proper clustering of various sequences (Figure3). Herein, the grouped sequences exhibit a high similarity in comparison to the other dataset entries, indicating a statistically significant evolutionary link (Hutson et al., 2001). The ADCL, BCAT, DATA, RATA and PUF sequences orderly show a diverse similarity score ranging from 19.42 – 37.97, 12.12 – 53.85, 13.74 - 44.72, 6.92 – 31.03 and 11.94 – 54.31, as shown earlier (Diebold et al., 2002; Pavkov-Keller et al., 2016).

**Figure 3:**
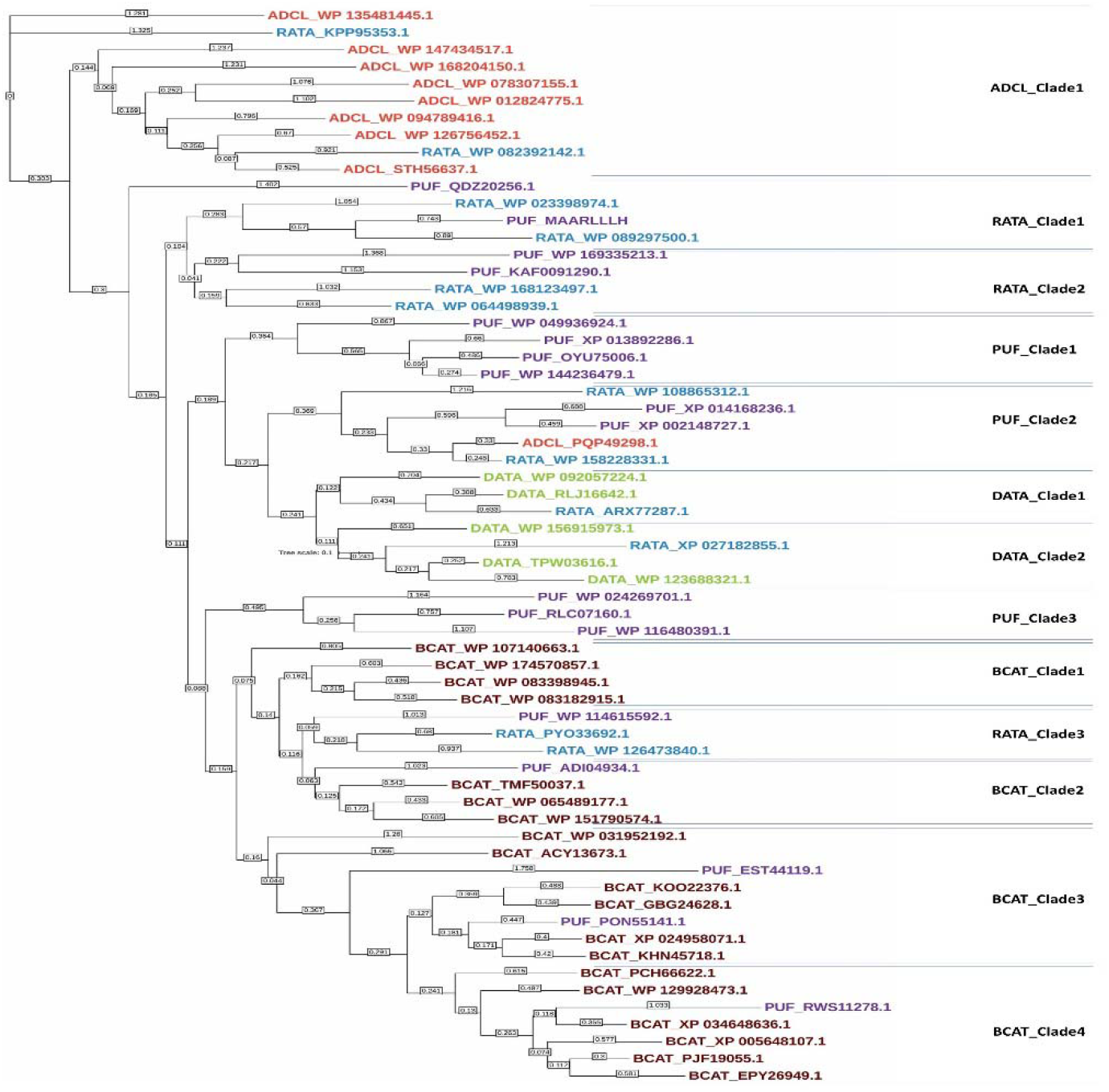
Phylogenetic analysis of the subfamilial datasets, indicating a segregated clustering and substantial diversity of several RATA and PUF entries.

Four BCAT, three RATA, three PUF, two DATA, and one ADCL clusters were identified, suggesting significant sequence divergence within the BCAT entries. The ADCL clade exhibited high branch length diversity, with a mean pairwise distance of 0.581 ± 0.468, ranging from 0.1 to 0.9 (O’Rourke et al. 2011; Porras Dominguez et al. 2024). Notably, an RATA entry RATA_KPP95353.1 shows a very high similarity of 96% to the closest ADCL sequence ADCL_WP_135481445.1. The two DATA clades show an average branch length score of 0.3903 ± 0.2576, unlike a bit higher score of 0.4338 ± 0.263, and 0.427 ± 0.1912, observed for the PUF-clades 1 and 2 (Jansonius, 1998). Contrary to this low average pairwise distances, the PUF-clade3 dataset displays a substantially higher branch length of 0.7558 ± 0.898, as observed for RATA clades 1 showing the respective score of 0.6658 ± 0.4136. Moreover, clades 2 and 3 of RATA show the respective score of 0.4338±0.4625, and 0.4735±0.406, unlike its clade3, indicating a substantial sequence diversification within the RATA sequences. Interestingly, RATA entries were found interspersed across other subfamilies, corroborating previous reports of their diverse phylogenetic affiliations (Hohne et al., 2010). Contrary to these subfamilial diversifications, BCAT clades indicated a very low average branch length score of 0.3467 ± 0.2543, and it goes in line with the earlier research (Maloney et al., 2010). These observations collectively suggest that RATA and PUF entries exhibit a broader phylogenetic distribution compared to BCAT and DATA entries, and is probably indicating that PUF dataset still encompasses the RATA sequences as well, and a need for a robust sequence sequence, viz. ADCL: STH56637.1 (*Escherichia coli*), BCAT: TMF50037.1 (*Chloroflexota bacterium*), DATA: WP_092057224.1 (*Planctomicrobium piriforme*), and RATA: WP_089297500.1 (*Actinoplanes -regularis*), is randomly selected and are modelled using Alphafold (Figure 4A) (Jhee et al., 2000, Burrell et al., 2010). To notably analyze the structural variations among these diverse sequences (Runthala & Chowdhury, 2016), their topologically conserved substructures, overlapped within 5□ distance deviation between the corresponding residues, are classification methodology (Runthala et al., 2024).

**Figure 4:**
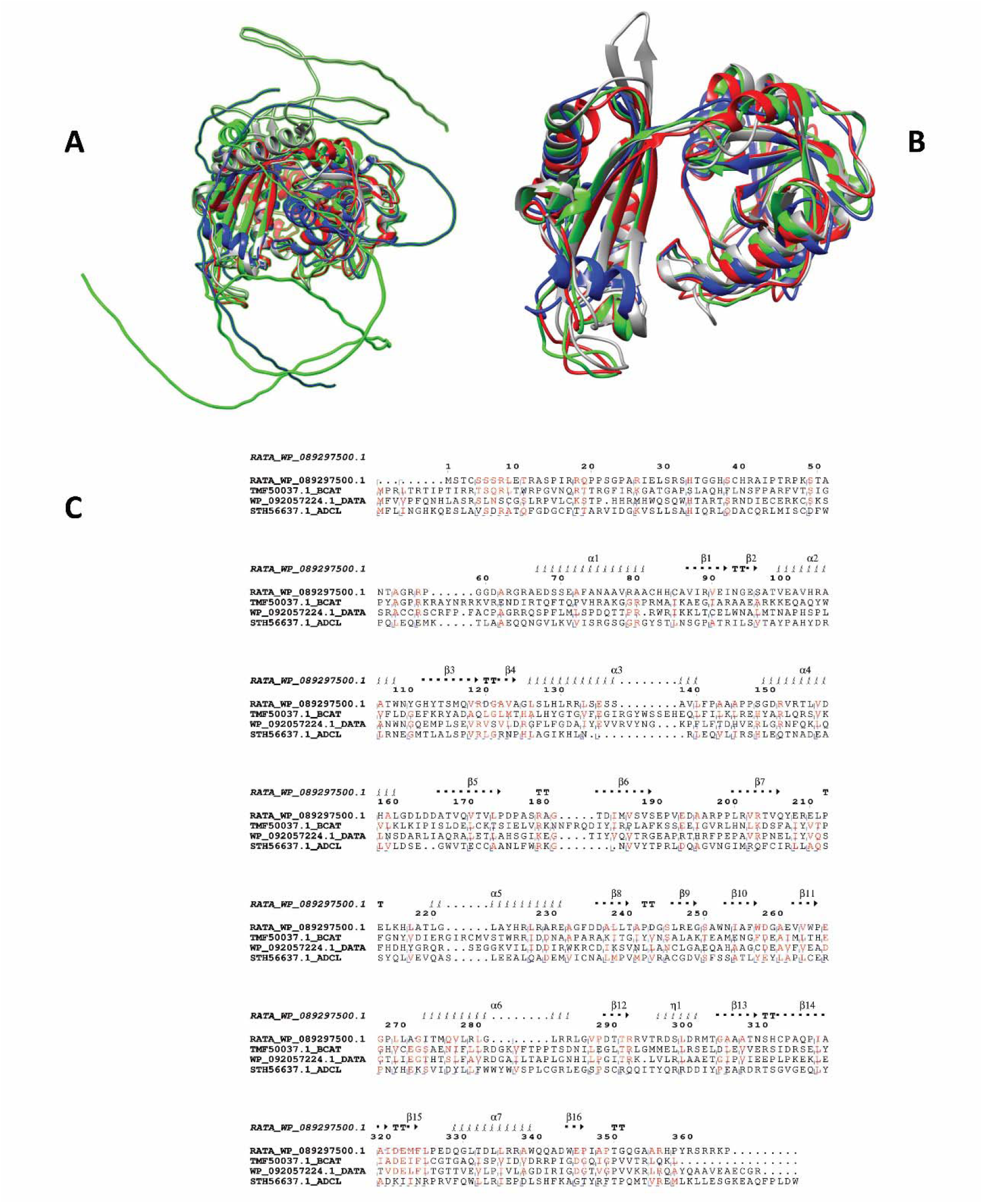
Superimposed Alphafold models of the (A) Complete protein models of the four subfamilial representatives STH56637.1(ADCL), TMF50037.1 (BCAT), WP_092057224.1 (DATA), and WP_089297500.1 (RATA), (B) Conserved core and (C) Sequence alignment, plotted as per the RATA representative, implying a remarkable structural similarity of the four subfamilies, despite a low sequence similarity.

For increasing the credibility of the downstream analysis, one longest subfamilial further analyzed using Chimera to map the functionally conserved structure, common across the four subfamilies (Figure 4B). Plotting the alignment conservation on the RATA representative WP_089297500.1 (Figure 4C) using Espript3 (Bonneau et al., 2009), it further confirms that the conservation is only pertained at a few functionally meaningful positions. Hence, despite the functional similarity, the transaminase family shares a strong structural similarity across the core (Pavkov-Keller et al., 2016).

### Analyzing the functionally crucial residues of the four subfamilies

The phylogenetic analysis unveiled the grouping of several subfamilial entries, although a low pairwise identity of 19.42%, 12.12%, 13.74%, 6.92%, and 11.94% is orderly observed among the ADCL, BCAT, DATA, RATA and PUF sequences. These findings suggest a high degree of evolutionary divergence within the transaminase family, potentially driven by the retention of only a handful of functionally critical residues. This observation underscores the limitations of contemporary motif-finding algorithms, which typically depend on substantial sequence homology for effective detection, especially for transaminases, as shown recently (Koper et al. 2022). Interestingly, these essential residues might even dictate the subfamilial affiliation, implying an intricate interplay between sequence variation and protein function.

To decode this further and refine the earlier benchmarked protocol (Figure 1), Weblogo (Crooks et al. 2004) is used for mapping the sequence conservation patterns to unveil the conserved positions, potentially pinpointing the functionally crucial residues that have withstood evolutionary pressures (Gavin et al., 2019). It results in the sequence logos where stacked symbols depict aligned residues, with symbol height reflecting amino acid frequency, normalized to 4 bits (Figure 5).

**Figure 5:**
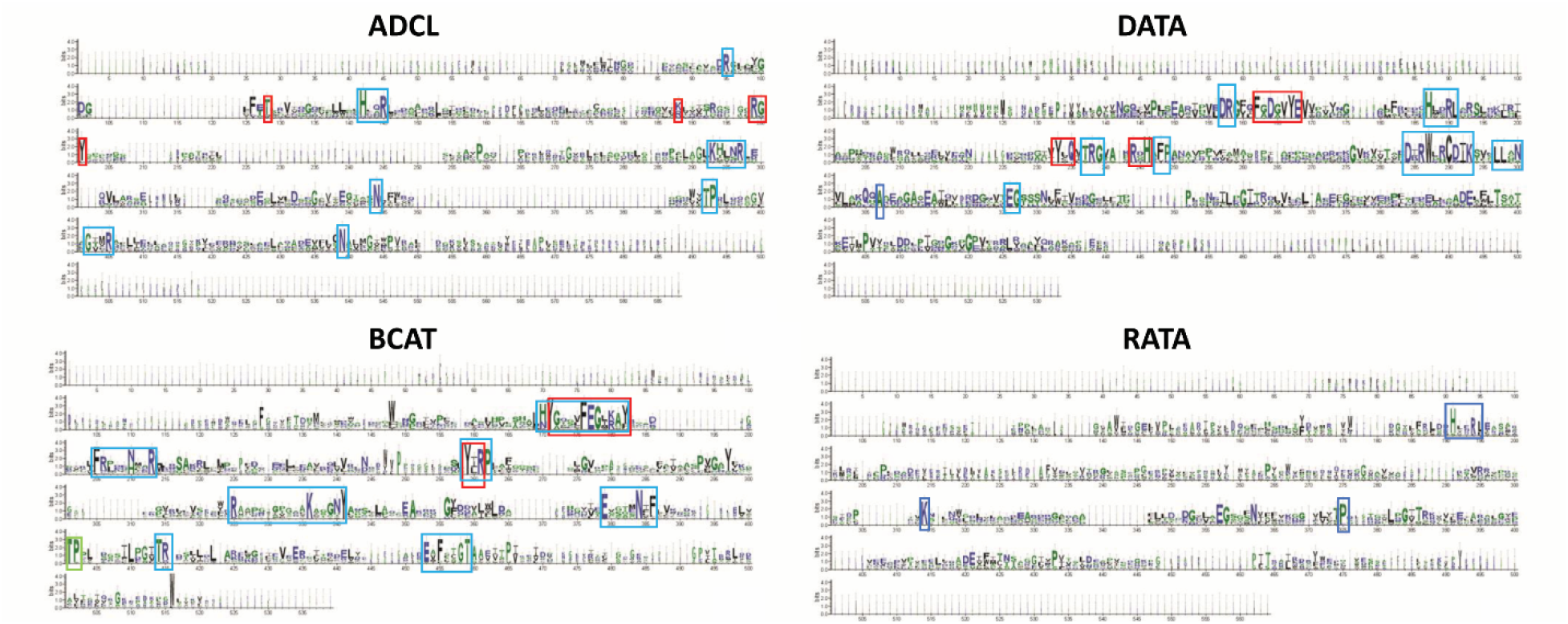
Weblogo representation of the sequence conservation for the four sequence datasets, identifying known motifs (red) and unveiling the novel ones (blue). It depicts the consensus sequence at each position, with stack heights representing the information content (bits) at that position. More prominent amino acids indicate higher conservation at that position.

Besides the already known motifs, the Weblogo analysis elucidates the novel conserved residue patterns for the four subfamilies ADCL, BCAT, DATA, and RATA. Surprisingly, it shows that despite the conservation of several other functionally crucial residues, the usually followed algorithm disregards them, highlighting its major limitation to robustly differentiate these subfamilies (Guan et al., 2015). Existing protocols define the ADCL subfamily based on an R or L motif near the RGY motif (positions 188 and 199-201 in our sequences), followed by a Threonine or Serine. Our sequence confirms this with an R at 188, RGY at 199-201, and a Threonine at 128. Interestingly, our analysis unveiled the novel conserved motifs (142-HXXR-145, 293-KXXXR-297, 392-TP-393, 402-GXXR-405) and individual amino acids (Asparagine at 344 and 449). These were identified within the secondary sheet structure of the sequence. These novel markers could potentially enhance future protocols for transaminase family identification. The presence of the YXQ motif is required to categorize a sequence to the DATA subfamily, and for our sequence, it is observed at the 233rd position, as demonstrated recently (Telzerow et al., 2021). Confirmatory motifs include the presence of two additional motifs: RxH at places 244–246, and FxxxxY (E, D) at positions 162-168. Interestingly, our analysis revealed seven novel conserved motifs (157-DR, 187-HxxRL, 237-TRG, 248-FP, 284-DxRWxxCDIK, 297-LLxN, and 326-EG), and these could prove to be crucial ones to reliably discriminate a sequence into DATA subfamily. These too had most of their motifs being linked into the secondary structure. The BCAT family is identified by the presence of 259-YxR-261 and 171-YxxxxFEGx-K/R-180 motifs (Zeifman et al., 2019). Additionally, the absence of K/R/H-182 residue further strengthens this assignment. Our analysis revealed novel conserved motifs within the sequence (184-Y/F, 210-N&H, etc.) and individual amino acid motifs (205-F, 213-R, 262-P) which were mostly attached to the sheeted portions and their presence would be valuable additions to future BCAT family identification protocols. 7 of 13 ADCL, 8 of 17 novel motif residues were found encoded within the secondary structure elements of these proteins, making them too crucial for the protein structure. Analyzing it further, it showed 1, 3, 2, and 1 novel conserved residue patterns of 293-KxxxR-297; 336-KxxxxY-341, 379-ExxxxNxF-386, and 453-ExFxxGT-459; 187-HxxRL-191, and 284-DxRWxxCDIK-293; and 191-HxxRL-195. Unlike other subfamilies, RATA family membership is confirmed by the absence of the RxH motif. Our sequence lacks this motif, supporting its classification as RATA. Notably, we identified novel sequence features within this family, including a conserved motif (191-HxxRL-195) and two conserved residues (K314 and P375) within the secondary structure elements. These findings could be instrumental in refining future RATA family identification protocols. Future studies should evaluate these novel motifs in the PUF sequences that remain unclassified using existing protocols.

## Conclusion

A comprehensive computational analysis used to identify and classify Transaminase enzymes by focusing on their subfamilies. Constructing a refined dataset of 247 transaminases from the initial dataset of 1035 proteins, and building their sub familial datasets ADCL, BCAT, DATA and RATA on the basis of known motifs, a substantial sequence diversity is observed within the subfamilial entries. Phylogenetic analysis further highlights the misclassification of RATA and PUF entries by clubbing entries within the tightly grouped other clades with a low branch length, and it refers to their incorrect functional assignment on the basis of known motifs. Further, as the conservation is only pertained to only the few meaningful sites across the four subfamilies, their functional motifs are reanalyzed through Weblogo, and it revealed novel conserved motifs for each subfamily. These computationally identified novel motifs will play a key role in the future studies regarding the subfamilial classification of transaminases, significantly enhancing the accuracy of allied experimental methodologies.

